# Engineering auronidin production in *Marchantia polymorpha* enables a new class of plant-derived textile pigments

**DOI:** 10.64898/2026.02.07.704544

**Authors:** Aurore Lormet, Jim Haseloff, Facundo Romani

## Abstract

Plant-derived pigments offer sustainable alternatives to synthetic colorants, yet their practical deployment in textiles is limited by restricted chemical diversity and low abundance. Liverworts represent a source of diverse chemical compounds, and the model liverworts *Marchantia polymorpha* is an emerging as chassis for bioengineering and synthetic biology. Here, we report the biotechnological application of auronidins, a rare class of flavonoid pigments, as textile dyes. Using the Marchantia, we engineered enhanced auronidin production through controlled expression of the R2R3-MYB transcription factor MpMYB14. We systematically benchmarked constitutive and inducible gene expression systems, including heat-shock, glucocorticoid receptor, and β-estradiol (XVE) circuits, identifying inducible strategies that decouple biomass accumulation from secondary metabolite production while achieving high pigment yields. Extracted auronidins were used to dye cotton yarn directly, demonstrating the feasibility of auronidins for textile dyeing. Our results establish Marchantia as a versatile plant chassis for programmable secondary metabolite production and introduce auronidins as a promising natural pigment platform for sustainable textile biotechnology.

## INTRODUCTION

Plant pigments are attracting attention as sustainable alternatives to synthetic colourants for foods, cosmetics and advanced materials. Besides their vivid colours, many plant-derived pigments offer antioxidant, antimicrobial, and anti-inflammatory activities that add value in functional foods and materials (Pizzicato et al., 2023). However, the palette of plant-derived pigments traditionally used for textile dyeing is chemically and chromatically constrained, being dominated by a limited number of compound classes such as anthraquinones, flavonoids, carotenoids, and indigoids. While these pigments have enabled natural dyeing practices for centuries, they occupy a restricted colour space and often display suboptimal stability or strong dependence on mordants and processing conditions. Moreover, many naturally occurring dyes accumulate at low concentrations or in highly specialized tissues, necessitating large-scale cultivation and extraction to achieve industrially relevant yields. These limitations represent a major bottleneck for the broader adoption of sustainable, plant-based dyes.

Plant metabolic engineering provides an efficient route to high levels of natural products with enhanced functional properties. Recent advances in crop engineering for pigment production show promise for sustainable textile applications (Ge et al., 2023). A biotechnological approach offers a route to overcome bottlenecks by enabling the controlled production of alternative pigment chemistries in engineered plant systems.

Liverwort species produce Riccionidin A, a unique type of red pigment that belong to the auronidin class (Berland et al., 2019). Notably, liverwort auronidins are predominantly associated with the cell wall rather than being vacuole-localised, distinguishing them from many commonly studied plant pigments (Jibran et al., 2024). In the model liverwort *Marchantia polymorpha*, the biosynthesis of auronidins is induced under environmental stress and nutrient deprivation and serves roles in photoprotection and pathogen resistance (Albert et al., 2018; Carella et al., 2019). This process is controlled by the R2R3-MYB transcription factor Mp*MYB14*, which is required and sufficient to activate endogenous biosynthetic genes (Albert et al., 2018; Kubo et al., 2018).

Marchantia polymorpha is an emergent chassis for plant synthetic biology (Sauret-Gueto et al., 2020). It has already been used for the production of proteins (Frangedakis et al., 2021; Tse et al., 2024b) and for the production of heterologous secondary metabolites (Forestier et al., 2025; Tse et al., 2024b; Zhang et al., 2020). A remaining challenge is the deployment of endogenous compounds with immediate functional applications that cannot readily be accessed in conventional chassis. Auronidin biosynthesis is a lineage-specific innovation of liverworts and provides an opportunity to extend the use of this system beyond proof-of-expression. Engineering auronidin accumulation therefore represents a logical progression in the development of Marchantia as a synthetic biology chassis. Here, we engineer plants with increased auronidin accumulation via MpMYB14 expression and report the first application of this class of natural products for textile dyeing. Together, these results highlight Marchantia as a promising biological chassis for metabolic engineering and auronidins as a distinct natural pigment for textile applications.

## RESULTS

Auronidins derive from the flavonoid pathway (Figure 1A) but the precise metabolic pathway for their biosynthesis in Marchantia has not been elucidated yet (Berland et al., 2019; Furudate et al., 2023). It has been previously shown that the transcription factor Mp*MYB14* is necessary and sufficient to induce the expression of enzymes associated with its biosynthesis and increase auronidin biosynthesis (Albert et al., 2018; Kubo et al., 2018). In plants with high accumulation of auronidins (_pro_*35S:*Mp*MYB14)*, analysis of visible absorption spectra of methanol extract from overexpressing plants showed an absorbance peak characteristic of auronidins at 490 nm, which has considerably lower intensity in the wild-type control extract and is absent in extracts of the Mp*myb14* loss-of-function mutant (Figure 1B). The peak coincides with the maximum absorption of the 4-neohesperidoside glycoside of the auronidin Riccionidin A (Berland et al., 2019). When excited at 490 nm, the extract displays a specific emission spectra maximum at 570 nm (Figure 1C). While absolute auronidin concentrations could not be determined due to the lack of purified standards, fluorescence (Ex 490 nm; Em 570 nm) provide a robust and specific comparative metric for subsequent analysis (Supplementary Figure 1).

**Figure 1.**
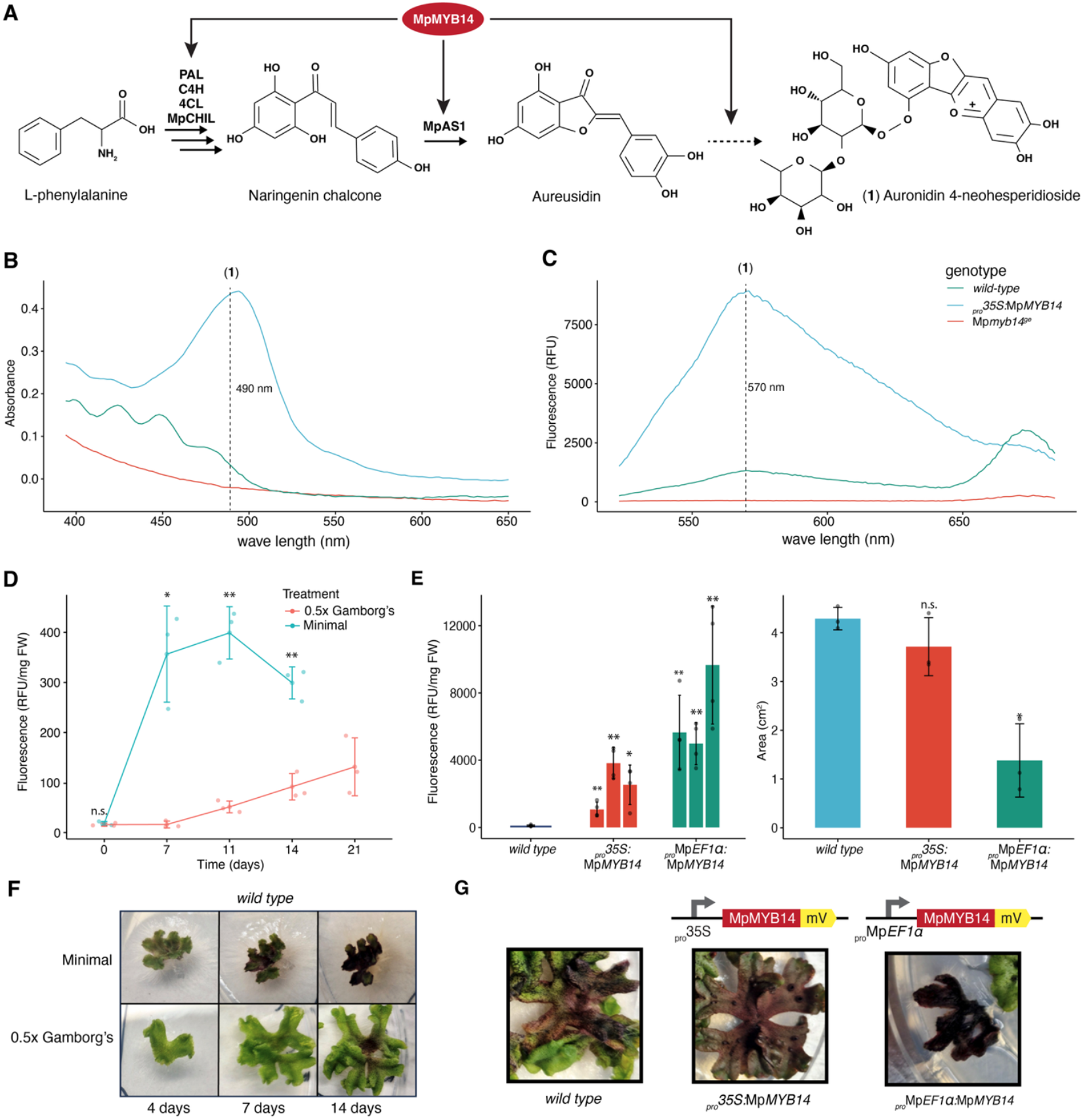
Biosynthesis of auronidins in Marchantia polymorpha. (A) Current pathway for auronidin biosynthesis in Marchantia. The dashed arrow indicate the reaction steps for which enzymes have not yet been identified. MpAS1, aureusidin synthase; MpCHIL, chalcone isomerase-like. PAL, phenylalanine ammonia-lyase; C4H, cinnamate 4-hydroxylase; 4CL, 4-coumarate: CoA ligase. Absorbance (B) and the fluorescence (C) spectra of methanol extracts from WT, myb14 mutant and _pro_35S:MpMYB14 Marchantia lines. The absorbance peak corresponding to 4-neohesperidoside auronidin (**1**) is highlighted as the main peak in the constitutive line (Absλ = 490 nm) and the corresponding emission peak (Emλ = 570 nm). (D) Time course of auronidin quantification in wild-type Marchantia polymorpha. Gemmalings were grown for 10 days under complete media (0.5x Gamborg’s B5 media) and either transferred to starvation conditions (1% sucrose (w/v) media) or maintained in complete media. Quantification of auronidins as accumulation of relative fluorescence units per milligram of fresh weight (RFU/mg FW) over time. (E) Representative pictures of plants under starvation (top) or control (bottom) at days 0, 4, 7, and 14. (F) Left, quantification of auronidins in 3-week-old M. polymorpha gemmalings overexpressing MpMYB14 under _pro_35S, wild type control, and mature sporelings of equivalent developmental time overexpressing MpMYB14 under _pro_MpEF1α. Three independent transgenic lines are shown. Right, thallus exposed area of 2-weeks-old sporelings from primary transformants. Sporelings were used as _pro_MpEF1α:MpMYB14 plants are unable to produce viable gemma. (G) Representative pictures of transgenic lines and wild type control. Error bars indicate means +/-SD of at least three biological replicates, individual values are shown; significant differences were tested using pairwise t-test comparing each time to the corresponding control and corrected by Bonferroni (* q-value < 0.05, ** q-value < 0.01).

Under standard *in vitro* culture conditions, auronidins are almost undetectable, with concentration estimated at around 14–30 RFU/mg FW (relative fluorescent units per milligrams of fresh weight). In stress, and particular nitrogen starvation conditions, plants accumulate auronidins in higher quantities (Zhou et al., 2024). To monitor the dynamics of auronidin accumulation, we grew wild-type plants for ten days before being transferred to either stress-inducing minimal media containing only 1% sucrose or complete media (0.5× Gamborg’s B5), as described before (Albert et al., 2018). In minimal media, auronidin concentration peaks at ∼400 RFU/mg FW after 11-days. Over time, adult plants on the complete medium also began to accumulate pigment in the centre of the thallus but in relatively lower quantities (Figure 1D). This was accompanied by strong growth defects in minimal media, consistent with previous reports (Berland et al., 2019). This provides an initial point to compare auronidin bioproduction and emphasises the importance of engineering strategies to increase production and mitigating growth penalties.

Increasing the levels of MpMYB14 represent a logical approach for engineering auronidins bioproduction. First, to benchmark different constitutive promoters and compare with natural production, we generated transgenic lines overexpressing Mp*MYB14* under the _pro_*35S* promoter against the stronger promoter _pro_Mp*EF1α* (Tse et al., 2024a). As expected, _pro_Mp*EF1α:*Mp*MYB14* lines achieved the highest levels of auronidin fluorescence (∼9,000 RFU/mg FW), followed by _pro_*35S:*Mp*MYB14* lines (∼4,000 RFU/mg FW). These values represent approximately 450-fold and 200-fold increases relative to unstressed wild type and 22-fold and 10-fold increases compared to the highest auronidin concentrations observed in nitrogen-stressed wild-type plants.

_pro_Mp*EF1α* plants exhibited uniform, intense red pigmentation across the thallus, whereas _pro_35S plants showed concentrated red coloration in the thallus centre with greener thallus margins (Figure 1E). The high auronidin levels in _pro_Mp*EF1α* lines was accompanied by a significantly smaller plant compared to wild type. On the other hand, the _pro_35S lines displayed a modest size reduction compared to wild type (Figure 1E). This trade-off between bioproduction and growth when transgenes are under different promoters was also described previously (Tse et al., 2024a).

To bypass the trade-off, inducible systems have been developed that allow decoupling the production of recombinant products from growth and allow tuneable expression (Liu et al., 2024). In Marchantia, the heat-shock inducible promoter _pro_Mp*HSP17*.*8A1* has been used to produce protein levels of betanins (Tse et al., 2024a). Alternatively, the glucocorticoid receptor (GR) system which regulates nuclear-cytoplasmic location (Ishizaki et al., 2015), and the β-estradiol inducible system are also compatible with Marchantia using (Ishida et al., 2022).

To compare different inducible strategy, we designed different combinations of genetic parts driving the inducible expression of MpMYB14 (Figure 2A). The transformed plants were initially grown under standard conditions for 10 days before being transferred to inductive conditions for 14 days and harvested. First, with _pro_MpHSP17.8A1 induction for 2 hours at 37 °C, we observed a modest increase in auronidin production and strong growth penalty, likely due to heat-induced stress (Figure 2B, Supplementary Figure 2). This highlights the limitations of transient induction for sustained metabolite accumulation, compared with continuous chemical induction via hormone-responsive systems. For the GR construct, we used the strongest promoter available 2×35S (Tse et al., 2024a) and observed a strong increase of up to ∼2,000 RFU/mg FW (Figure 2C). Alternatively, the _*pro*_Mp*E2F:XVE >>* Mp*MYB14* synthetic circuit produced up to ∼2,800 RFU/mg FW, suggesting that the XVE system is a powerful tool to drive high levels of expression with tight regulation (Figure 2D). This is particularly due to the amplification step of the XVE system (Figure 2A). Yet, the _*pro*_Mp*E2F* is a weak promoter compared to others and is only active in dividing cells (Romani et al., 2024). To obtain even higher levels, we built an alternative version replacing _*pro*_Mp*E2F* with _*pro*_*35S*. Overall, _*pro*_*35S:XVE >>* Mp*MYB14* resulted in even higher yields, with up to ∼5,000 RFU/mg FW but also showed significant activity under non-induced conditions (Figure 2E). Overall, chemical induction of Mp*MYB14* could achieve yields comparable with the best constitutive promoters. Under our regimen, GR and _*pro*_Mp*E2F:XVE* did not compromise the initial vegetative growth phase under the conditions tested (Supplementary Figure 2).

**Figure 2.**
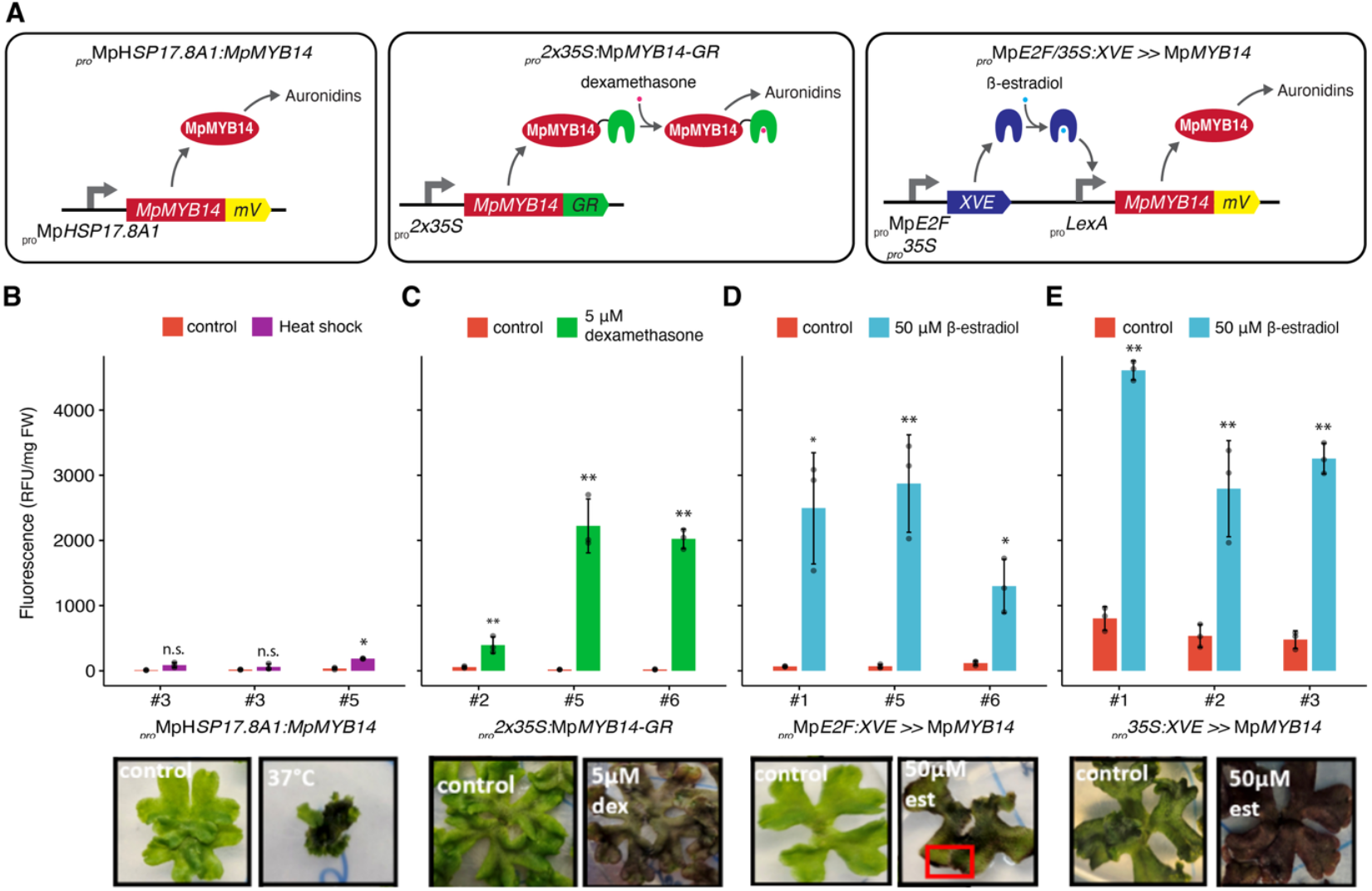
Inducible expression of auronidin. (A) Design of gene construct for auronidin expression(B-E) Quantification of auronidins in M. polymorpha gemmalings from inducible lines (_pro_MpHSP17.8A1:MpMYB14, _pro_35S:XVE >> MpMYB14, _pro_MpE2F:XVE >> MpMYB14, _pro_35S×2:MpMYB14-GR). Plants were grown for 10 days in standard conditions and then exposed to two rounds of heat-shock (37 °C) for two hours per day or transferred to inducible media as indicated in each plot (F-H). Control plants were either not exposed to heat-shock or transferred to media without inducer present. Auronidin levels were quantified after 14 days. Three biological replicates and three independent transgenic lines were used. Bottom, representative pictures of transgenic lines and their respective control. Error bars indicate means +/-SD of at least three biological replicates, individual values are shown; significant differences were tested using pairwise t-test comparing each time to the corresponding control and corrected by Bonferroni (* q-value < 0.05, ** q-value < 0.01).

To assess the potential of engineered auronidin production for direct applications, we scale-up the _*pro*_*35S:*Mp*MYB14* transgenic line to obtain higher biomass by growing plants in hydroponics for three weeks. The resulting extract was filtered and directly used to dye cotton yarn using an exhaustion dyeing protocol at 30% weight of fibre, without the use of mordants.

After dyeing, the cotton yarn exhibited a robust pink coloration under natural light compared with yarn dyed using wild-type extracts (Figure 3A–B). In addition, dyed yarn retained strong auronidin-associated fluorescence (Figure 3B). We next evaluated the effect of commercially available pre-mordant treatments and tested wash fastness. Yarn pre-treated with tannins displayed increased colour intensity and improved retention after washing compared with unmordanted controls (Figure 3C). Yarn pre-treated with tannin and aluminium acetate showed a further increase in colour intensity and minimal visible colour loss after washing in 50 °C water. Together, these results provide initial evidence that auronidins from engineered Marchantia can be applied as functional textile pigments, with colour depth, fluorescence and and reasonable fastness properties within the range observed for commonly used natural dyes.

**Figure 3.**
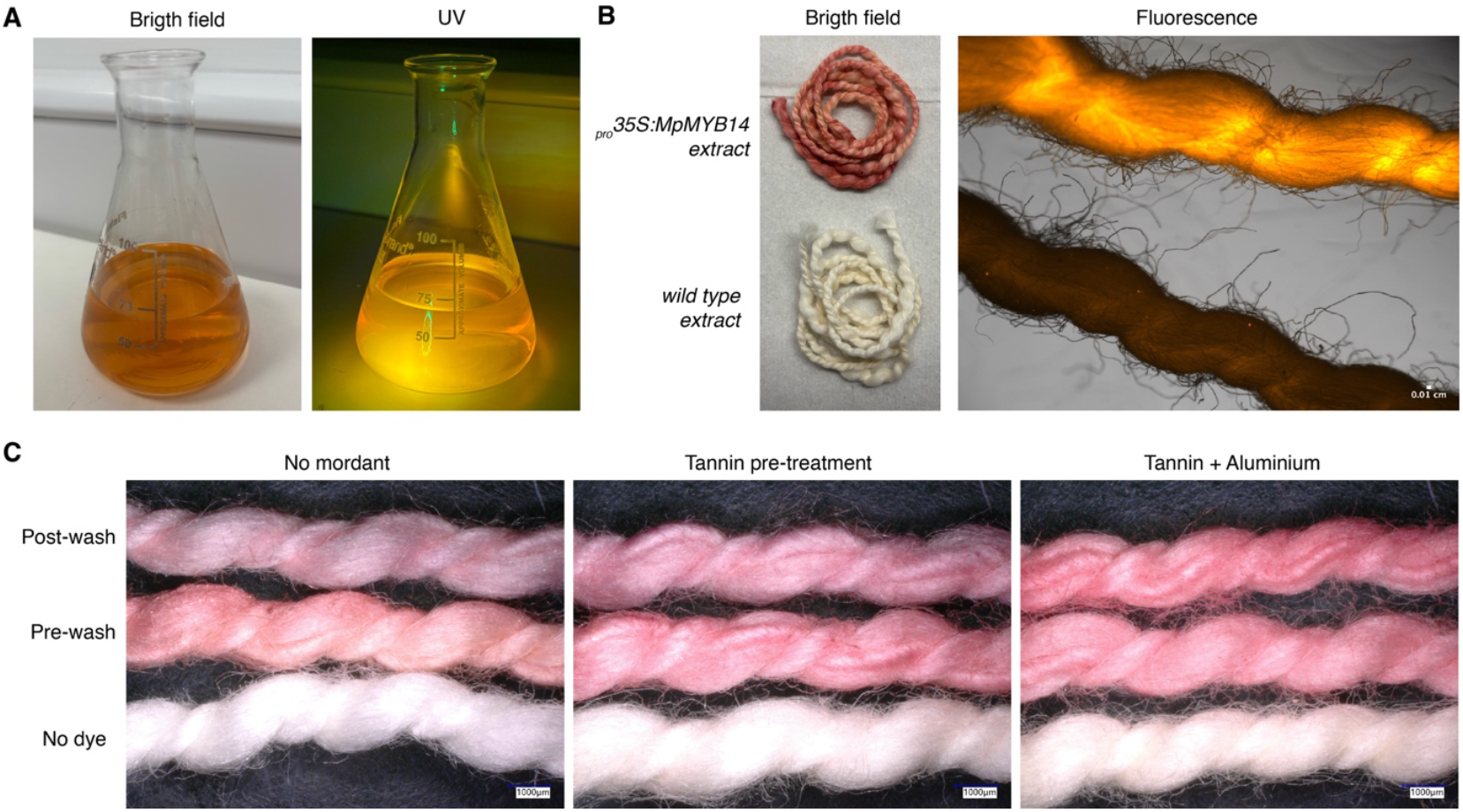
Auronidin extract as textile dyes. (A) A methonal extract from a scaled up culture of _pro_35S:MYB14 thallus in bright light conditions and under 365 nm UV light with a ∼400 nm bandpass filter. (B) Cotton fibres dyed with auronidin pigment extract from Marchantia _pro_35S:MYB14 line or wild-type plants. Microscopy images under the bright field or fluorescence images of the same fibres under a YFP band pass filter (500 +/-20 nm excitation and 535 +/-20 nm emission) and coloured using the Orange Hot LUT. (C) Representative images of dyed yarn without mordants or pre-treated with tannins (gallnut) or aluminium acetate. Scale bars are shown in the respective picutres.

## DISCUSSION

Here, we exploited Marchantia to engineer substantially elevated levels of auronidins. Compared to wild-type strain, we observed up to >100-fold higher auronidin yields. MpMYB14 overexpression functions as an effective lever to upregulate native biosynthetic genes and increase auronidin production. These features, together with the capacity for simple cultivation and fast growth of Marchantia, highlight its wider potential as a green biofactory for secondary metabolism. As shown before, constitutive bioproduction is accompanied by a trade-off in growth (Tse et al., 2024). Here, we further compare the XVE and GR inducible circuits that have the capacity to mitigate fitness costs by temporally separating biomass accumulation and secondary metabolism. The XVE system under the _*pro*_Mp*E2F* exhibited the best dynamic range among the tested conditions, and the new _*pro*_*35S* version can provide higher yields but with leaky expression.

Auronidins have not previously been explored as textile dyes, and our dyeing experiments establish a direct biotechnological application for this class of plant pigments. By introducing auronidins as functional textile pigments, this work expands the repertoire of plant-derived dyes by adding a pigment class not previously exploited. In addition to textile dyeing, their intrinsic properties (including fluorescence, UV absorption, and reported antimicrobial activity) may be relevant for further expanding their range of applications.

Recent work has shown that auronidins can undergo polymerisation and accumulate as insoluble complexes associated with plant cell walls (Jibran et al., 2024), revealing a level of chemical complexity that distinguishes them from many commonly used natural dyes. Dyeing activity in our experiments is attributable to the soluble auronidin fraction, highlighting polymerisation state as a key variable for future optimisation of extractability and dyeing performance.

The bioproduction of auronidins is a powerful proof-of-concept demonstrating the unique advantage of producing endogenous chemicals that cannot be readily produced in conventional chassis. Beyond Marchantia, liverworts possess a rich biochemical diversity that can be unlocked. Marchantia offers a closely related chassis with shared precursors and metabolism, which has the potential to enhance the accessibility of liverwort-specific metabolites. A remarkable example is perrottetinene, a psychoactive cannabinoid-like compound present in the leafy liverwort *Radula marginata* (Chicca et al., 2018), along with other bis-bibenzyls and terpenoids with potential medicinal applications (Sen et al., 2023). Together, these results position Marchantia as a versatile platform for translating lineage-specific plant chemistry into functional applications

## ACKNOWLEDGEMENTS

F.R. is a Leverhulme Early Career Fellow (ECF-2023-534) funded by the Leverhulme Trust and the Isaac Newton Trust (23.08(f)). We would like to thank Rory Harrison for his help with hydroponics.

## AUTHOR CONTRIBUTIONS

FR conceived, designed and supervised the research. AL performed the experiments. FR and AL analysed and interpreted the data and wrote the manuscript. JH contributed to experimental design discussions and provided editorial feedback on the manuscript.

## MATERIALS AND METHODS

### Plasmid assembly

Constructs were obtained from the one step GoldenGate cloning protocol for *Marchantia* gene cassette described in (Tse et al., 2024a) using the pBy10 acceptor. L0 parts from the OpenPlant Toolkit (Sauret-Gueto et al., 2020) were used as specified in Supplemental Table 1. The coding sequence of MpMYB14 (Mp5g19050) CDS12 part was described in (Kongsted et al., 2025). The presence of the correct insert was confirmed by restriction digestion and Sanger sequencing.

### Agrobacterium-mediated transformation

*Marchantia* spores were sterilised as previously described (Sauret-Güeto et al. 2020) and grown on solid 0.5x Gamborg B5 medium for 5 days. A modification of the original *Marchantia polymorpha* sporeling Agrobacterium-mediated stable transformation method was used to transform the *Marchantia* cam-1/2 spores as described before (Annese et al., 2025).

### Plant materials and growth conditions

*Marchantia polymorpha subs. rudelaris* accessions Cam-1 (male) as the wild-type and Cam-1 x Cam-2 spores for transformation were used. The Mp*myb14* knock-out in the Cam background was described in Kongsted et al. (2025). Under standard conditions, lines were grown on complete media containing 0.5x Gamborg B5 medium with 1.2 % (w/v) agar micropropagation grade (Phytotech #A296) at pH 5.5 and grown under continuous light at 21°C with a light intensity of 150 μmol m^-2^ s^-1^.

For nutrient deprivation experiments (9 plants per plate / biological replicate; 3 biological replicates per treatment), gemmae were plated on 0.5x Gamborg media, as described before (complete media) and grown for 10 days under standard growing conditions. After that, for starvation treatment (Albert et al. 2018), plants were transferred to minimal media containing 1 % (w/v) sucrose, 1.2 % (w/v) agar at pH 5.8 and further grown under standard conditions. Alternatively, controls plants were transferred to complete media again.

For experiments with inducible expression (three gemmae per line; three lines per independent transformant), gemmae were plated on complete media and grown under standard conditions for 10 days. After this time, transformants were transferred to complete media without supplementes or supplemented with 5 μM of dexamethasone (#265005, Sigma-Aldrich) or 50 μM of 17-β-estradiol (#E8875, Sigma-Aldrich). Plants were grown on inducible media for 14 days before harvest. Similarly, transformed plants with the heat shock promoter were grown for 10 days in standard conditions and placed in a 37 °C incubator without light for 2h a day for 2 days before being moved back to the standard growth conditions for one week.

For scaling up, the *Marchantia* line plants were placed in hydroponic vessels allowing a continuous flow of nutrient film solution: 12 mL each of Samurai Hydro Grow Nutrient A and B (Shogun fertilisers) in 4 L of tap water. The pieces of meristems cut from plants were anchored on a “mat” and covered with a propagator lid, creating a humid environment, over which an LED panel was mounted for optimal growth. Each *Marchantia* line was grown for 3 weeks before induction and extraction of auronidin.

### Auronidin extraction and quantification

The pigment was extracted from fresh thallus tissue using methanol: water : formic acid (80:19:1) as described before (Albert et al., 2018). All analyses were done using at least 3 replicates. Approximately 100 mg of thalli were crushed using a plastic pestle in a 1.5 mL tube in the presence of 1 mL of the solvent (at 1:10 material-liquor ratio), vortexed for 10 seconds, and centrifuged for 2 minutes at 13,300 rpm. 50 μL of the supernatant were transferred to a 384-well polypropylene microplate (Greiner #781209) to be analysed in a BMG ClariostarPlus plate reader. For fluorescence quantification, excitation at 490-15 nm and emission 570-8 nm (gain: 1512) using the fluorescence intensity end-point mode and normalised with respect to fresh weight. Sample values were collected as 1 mm orbital average and adjusted by subtracting the fluorescence values of the blank. To optimise the emission and excitation parameters we followed the literature (Berland et al., 2019) and measured the absorbance spectra (350-650 nm) using the absorbance mode, and the emission using the fluorescence intensity emission spectrum mode, collecting from 549-660 nm with excitation at 490-15 nm. Serial dilutions were performed to confirm the linear relation of the signal (Supplemental Figure 1).

### Dyeing fibres

Cotton fibres were dyed using an auronidin *Marchantia* pigment extract by a methanol solution as described before but using a material-liquor ratio of 1:4. We adapted a protocol for exhaustion dyeing method described before for Sorghum husk (Hou et al., 2017). Briefly, commercial cotton yarn was first scoured in water with soup for 40 minutes and then incubated with the extract at 30% weight of fibre ratio at room temperature until the solvent was completely evaporated overnight. Gallnut and aluminium acetate (Wonkyweaver) following manufacturing instruction at 10% weight of fibre ratio. Fastness was tested using distilled wated at 50 °C for 30 minutes and dried.

### Data analysis

To obtain plots and statistical analysis, data was processed using R v 4.4.1 software and RStudio. Statistical analyses were done using the built-in statistics package using pairwise t-test comparing each time to the corresponding control and corrected by Bonferroni using default parameters. Plots were made using the ggpubr package. Pictures were taken using either a Keyence VHX-5000 digital microscope or a Leica M205 FA stereo microscope equipped with a YFP fluorescence filter and analysed using ImageJ/Fiji. Thallus area, the exposed area was quantified from pictures with ImageJ/Fiji software with 3 biological replicates.

## SUPPLEMENTAL FIGURES

**Supplemental Figure 1.**
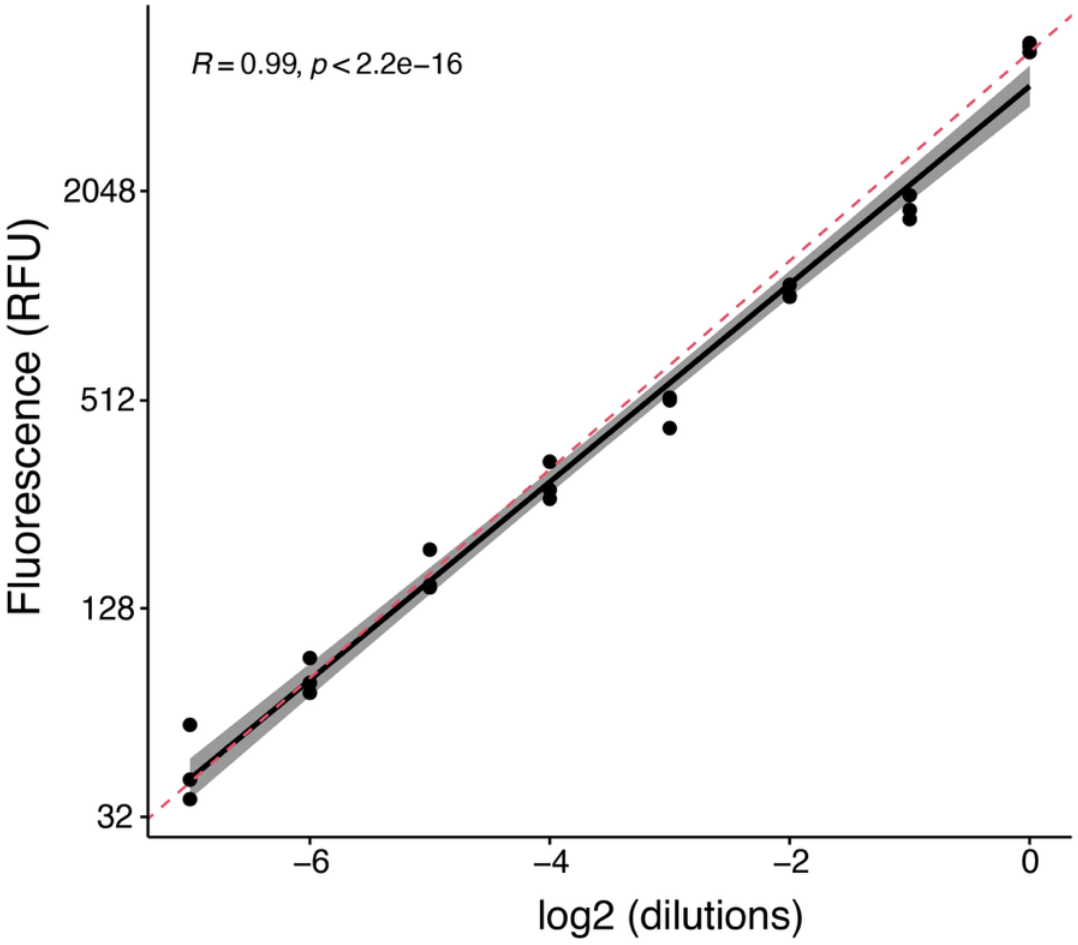
Serial dilution regression analysis shows linear fluorescence relationship.

**Supplemental Figure 2.**
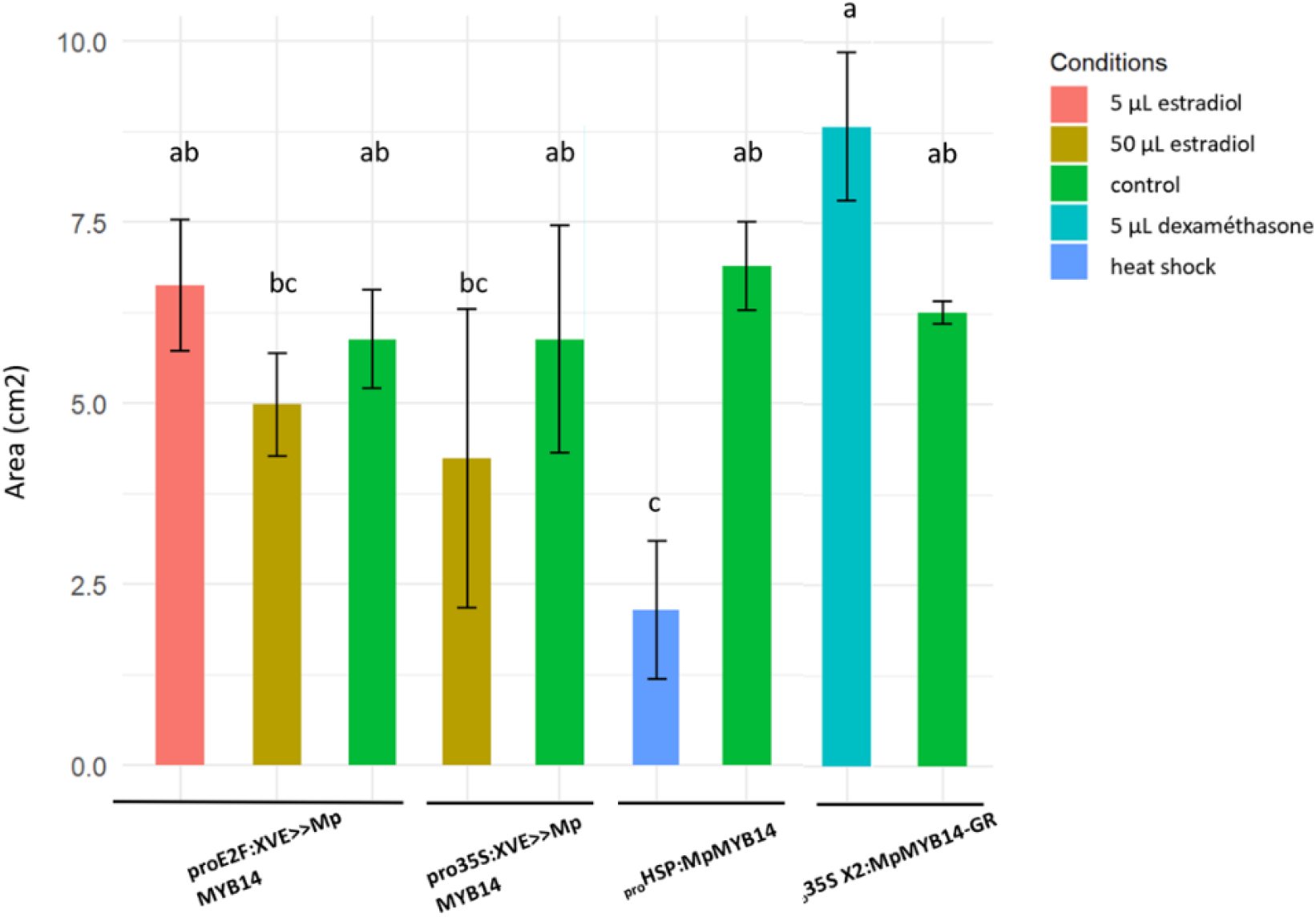
Relative thallus size of the 5 different inducible line plants (14 days old). Each bar represents an independent transformant for that promoter construct and the mean of three biological replicates with an error bar for standard deviation between the replicates. Letters above the bars indicate statistically significant differences between means (Anova and Kruskal test at P<0,05 respectively).

**Supplementary Table 1.** Vectors and parts used for cloning.

